# Age-related myelin deficits in the auditory brain stem contribute to cocktail-party deficits

**DOI:** 10.1101/2024.07.29.605710

**Authors:** Shani Poleg, Ben-Zheng Li, Matthew Sergison, Matthew Ridenour, Ethan G. Hughes, Daniel Tollin, Achim Klug

## Abstract

Age-related hearing loss consists of both peripheral and central components and is an increasing global health concern. While peripheral hearing loss is well understood, central hearing loss— age-related changes in the central auditory pathways resulting in a listener’s inability to process sound correctly —remains poorly understood. In this study, we focus on the pathway from the cochlear nucleus to the medial nucleus of the trapezoid body (MNTB), which depends on heavily myelinated axons for microsecond-level temporal precision required for sound localization.

Using a combination of auditory brainstem response recordings (ABR), advanced light and electron microscopy, and behavioral testing with prepulse inhibition of the acoustic startle response (PPI) we identified a correlation between oligodendrocyte loss, abnormal myelination in MNTB afferents, altered ABR wave III morphology indicating MNTB dysfunction, and deficits in spatial hearing behaviors in aging Mongolian gerbils.

These findings provide a mechanistic explanation of how demyelination contributes to age-related dysfunction in the auditory brainstem’s sound localization pathway.

## INTRODUCTION

Age-related hearing loss, or presbycusis, is a global health problem which is becoming increasingly important as many modern societies have progressively larger portions of aging members. Presbycusis affects approximately one-third of adults beginning at middle age, with over half of adults aged 70 and older experiencing it^1–4^. Additionally, untreated hearing loss has a significant impact on the quality of life, because in addition to the direct impact of the hearing loss itself, it is linked to an elevated risk of secondary conditions such as social isolation, depression, dementia, Alzheimer’s disease and many other mental health problems^2,5–9^. Thus, early detection and treatment of hearing loss is crucial for promoting healthy aging.

Peripheral hearing loss – the loss of hair cells in the inner ear – is one mechanism which contributes to presbycusis. As hair cells die during the lifetime of a human listener or other mammalian species, the thresholds for the detection of sound increase. This form of hearing loss is well understood and common treatments such as hearing aids or cochlear implants are effective to restoring audibility in most cases^10^.

By contrast, a completely independent mechanism of age-related hearing loss, here termed “central hearing loss”, is mediated by changes of neural circuits in the CNS, here termed “central hearing loss”. In contrast to peripheral hearing loss, central hearing loss is much less well understood and no treatments are available to date. While changes in several locations along the ascending auditory pathway have been postulated to contribute to central hearing loss, one major component involves the age-related reduction in binaural hearing abilities, such as listening to and effectively segregating speech in noisy environments in listeners who may have otherwise normal hearing thresholds^11–14^.

The ability to listen in complex multisource environments as well as the ability to pinpoint the source of a sound relies on a precise comparison of inputs from the two ears^15^. Extreme temporal precision is required because brainstem circuits must be able to detect differences in arrival times of sound waves between the two ears which can be on the order of just a few microseconds, thus two orders of magnitude faster than a typical action potential ^16^. To accomplish such high levels of precision, this process depends on a number of specializations along the sound localization pathway, one of which is fast synaptic transmission along heavily myelinated axons in the brainstem^16–18^. The medial nucleus of the trapezoid body (MNTB), which provides fast and temporally precise glycinergic inhibition to the principal binaural nuclei, the lateral and medial superior olive (LSO and MSO, respectively), stands out for having both one of the largest synapses in the brain (the calyx of Held) and wide-diameter afferent axons, underscoring the critical need for temporal precision in this circuit^18–22^. As a result of these specializations, MNTB neurons exhibit phase locking, a measure of the temporal fidelity of spike timing relative to the ongoing phase of the sound stimuli, that exceeds that observed in auditory nerve fibers^23,24^.

Due to the high level of adaptation aimed at preserving temporal fidelity that is built into this system, it seems intuitive that even minor disruptions in the temporal precision in this pathway can lead to significant deficiencies in sound localization. Alterations in myelination within the auditory brainstem affect the speed of electrical propagation and, consequently, the timing of auditory brainstem activity^18,20,25–27^. Evidence from various studies indicates that myelination deficiencies contribute to the aging brain phenotype^28,29^, begging the question whether age-related myelination deficits might contribute to a decline in timing within the binaural pathway, which in turn might contribute to central hearing loss. Binaural hearing is known to mediate not only sound localization abilities per se but also help a listener in acoustically busy environments such as crowded restaurants where multiple sound sources are active simultaneously and a listener wants to focus on a single sound source such as the voice of a conversation partner. In such situations, often termed “cocktail party situations”, the sound localization circuit helps listeners to isolate the sound source of interest from distracting background noises based on spatial location^30–32^. Thus, a listener affected by age-related decline in spatial hearing should primarily notice a decline in their abilities to focus on a sound of interest within background noises^14^.

The decline in temporal processing associated with aging is not unique to humans; rather, it appears to be a widespread phenomenon among mammals^33,34^. The Mongolian gerbil (*Meriones unguiculatus*) is often used as an animal model for human hearing, including age-related alterations in the sound localization pathway. Gerbils possess distinctive features such as well-developed low-frequency hearing and low frequency sound localization abilities^35^ while exhibiting minimal age-related loss of hair cells in the inner ear^36,37^. This renders them better models for studying human hearing compared to other rodents, such as mice and rats^38,39^.

In our study, we employ a combination of behavioral and electrophysiological assessments of auditory circuits, immunohistochemistry, electron microscopy (EM) and coherent anti-stokes Raman spectroscopy (CARS) to investigate the contribution of the myelin microstructure to sound localization in aging gerbils compared to young adult controls^40,41^. We found that gerbils recapitulate the age-related binaural hearing deficits that are so common in humans. We assessed spatial sound discrimination abilities using a behavioral paradigm measuring prepulse inhibition of the acoustic startle response (PPI). PPI is a reflex that occurs in most mammals, including gerbils ^42–44^. Rodents will startle when presented with a loud, brief stimulus, but will reduce the amplitude of their startle response if a detectable change in the sensory environment, or prepulse, occurs just prior to the startle sound presentation ^42,45^. Because PPI is a reflex and requires no training, PPI offers a high throughput method to test spatial hearing capabilities in a large cohort of animals ^43–45^. The brainstem circuit that encodes PPI is well researched and includes the binaural brainstem nuclei that initially encode the acoustical cues to sound location ^46^, thus allowing the use spatial cues as the prepulse. With a spatial speaker swap task acting as the prepulse cue, we found that aging animals have lower levels of PPI in response to wider angles than young animals, indicating an impairment of spatial hearing abilities.

In both gerbils and humans these spatial hearing deficits can be measured noninvasively as significantly attenuated wave-III and earlier onset wave-IV in click ABRs from old gerbils, suggesting degradation of inhibitory inputs and reduced MNTB activity in the superior olivary complex. Aging gerbils also exhibited reduced amplitudes of their binaural interaction component (BIC) of ABRs^47^, which is a measure of the efficacy of binaural processing in the sound localization pathway of small animals.

The neural mechanism that mediated both the behavioral and physiological changes is an age-related demyelination of afferent fibers to the MNTB. In aging gerbils, these fibers exhibited significantly reduced myelin thicknesses and axon diameters, leading to suboptimal action potential transduction. To uncover potential mechanisms underlying alterations in myelin structure, we investigated the numbers of oligodendrocytes precursor cells (OPCs) and mature, myelinating oligodendrocytes (OLs). Our results demonstrate a deficit in OL numbers in old gerbils, without a change in the total number of oligodendrocyte precursor cells, suggesting a potential mechanism by which auditory phenotypes might arise in aging.

## RESULTS

The goal of this study was to test the overarching hypothesis that age-related alterations in the binaural circuits in the auditory brainstem contribute to a decreased performance in binaural hearing tasks. This decreased performance in turn results in impaired spatial hearing and impaired cocktail party performance. We used Mongolian gerbils as our model, with groups of young (2-3 months old) and aged animals (2.5 to 3.5 years old). All experiments were conducted on the same cohort of animals to determine whether physiological responses correlated with histological findings.

### Aging gerbils show compromised spatial hearing

We used paired pulse inhibition (PPI) as a behavioral assay to assess the spatial hearing of young and aging gerbils, based on methods previously used in a Guinea Pig model in the lab^48^. This behavioral test requires no training of the animals but rather relies on the fact that mammal including humans startle when a loud sound is presented unexpectedly (Figure 1A and 1C), and the amount of startle can be quantitatively measured (Figure 1B). Whenever something changes in the animal’s sensory environment that is detectable just before the startle sound, a midbrain reflex circuit (partially) inhibits the startle in a graded way that scales with detectability of the change. We made use of this reflex by introducing gaps into ongoing background noise (Figure 1E) or by changing the location from which test sounds were emitted (Figure 1A and 1G).

**Figure 1.**
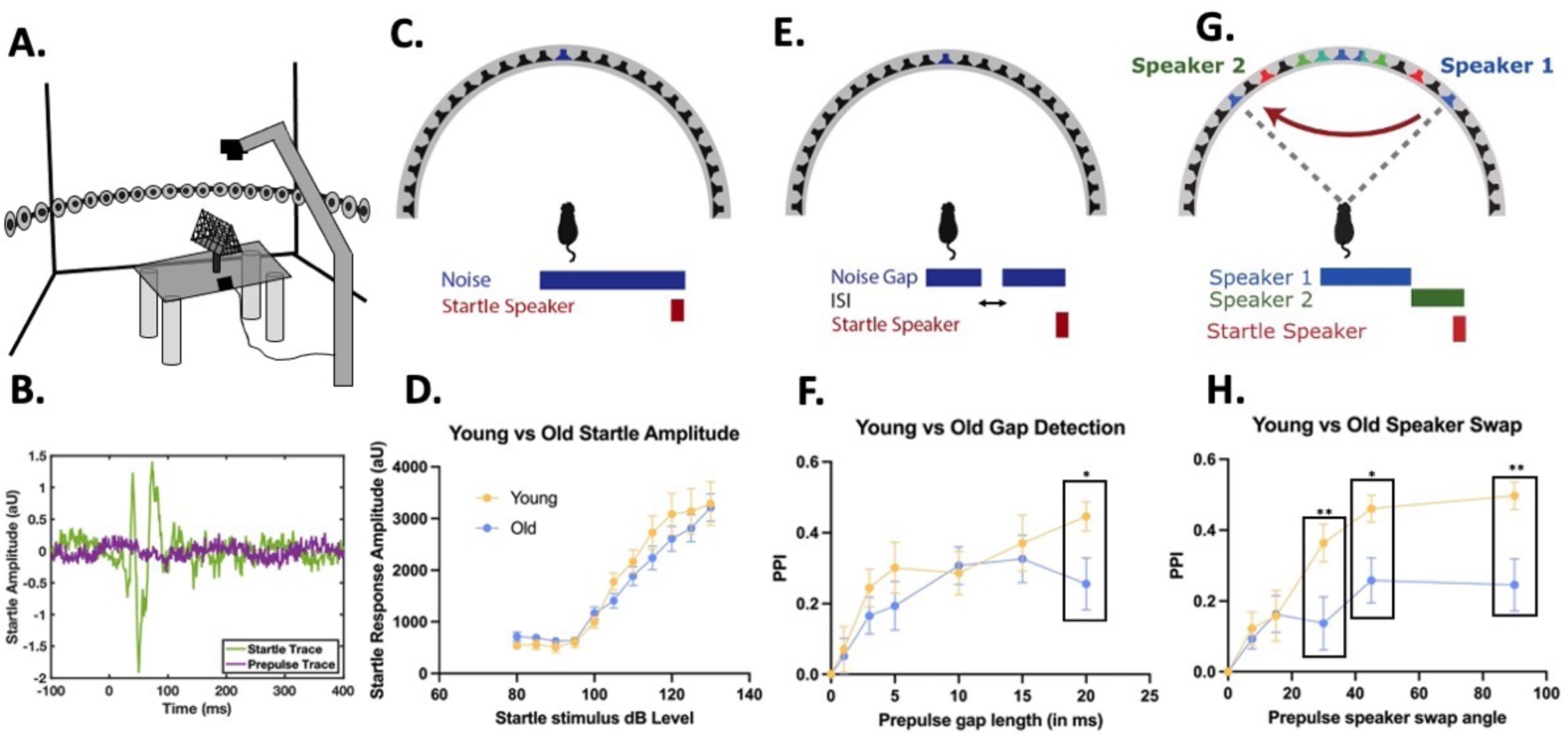
Prepulse inhibition of the acoustic startle response reveals spatial hearing deficits in aged gerbils. **(A)** Diagram of acoustic startle setup. Prepulse stimuli are presented from a 24 speaker array located at midline. Startle stimuli are presented from a startle speaker above the animal. Animals are held at midline while being tested. Startle response is measured on an accelerometer attached to the animal’s cage. **(B)** Example startle trace showing measured startle response from a startle (green) trial and a prepulse plus startle (purple) trial, demonstrating prepulse inhibition of the acoustic startle. **(C)** Schematic showing startle amplitude response testing. Startle stimuli were presented at 5 dB increments from 80dB to 130 dB. **(D)** Startle amplitude response in arbitrary units to varying dB levels of startle. No difference between aged and young animals was found. Young n=10, old n=21. **(E)** Schematic of gap detection task. A gap of variable length was presented in broadband noise 80 ms prior to startle. **(F)** PPI value as a result of gap length. Squares outline conditions with significant difference in PPI values between young and old animals. Young n=10, old n=20. **(G)** Schematic of speaker swap task. Broadband noise was swapped from a speaker on one midline of the animal to another speaker across the midline at 80 ms prior to startle. **(H)** PPI value as a result of speaker swap angle. Mean and standard error of the mean are plotted for all groups. Diagrams modified from Greene et al. 2018^44^.

First, we assessed the ability of each young and aging gerbil to elicit a startle response to a startle stimulus presented by itself (Figure 1B). To this end, we presented brief, broadband noise pulses as startle stimuli with varied intensities from 80 to 130 dB SPL in 5dB increments (Figure 1C). Startle amplitudes of the animals varied with the sound intensity of the startle sound (F=46.10). We found that in response to 105 dB stimuli or louder, startle amplitudes began to increase from baseline levels in young animals (105 dB: p=0.0396, 110 dB: p=0.0006, 115 dB: p<0.0001, 120 dB: p<0.0001, 125 dB: p<0.0001, 130 dB: p<0.0001). In the older animals, startle amplitudes began to increase above baseline at sound levels of 110 dB (p=0.0002) (115 dB: p<0.0001, 120 dB: p<0.0001, 125 dB: p<0.0001, 130 dB: p<0.0001). We selected a startle stimulus level of 117 dB based on these results, as this elicited a large enough startle response compared to baseline that could then be modulated by a prepulse stimulus. There was no variation in startle amplitude as a result of age (F=1.781) and no interaction between age and stimulus level effecting startle amplitude (F=0.7245), indicating that impairments seen later in our aging animals are not due to an inability to startle.

Next, we presented a gap in 70 dB broadband noise at varying lengths to act as a prepulse prior to the startle stimulus (Figure 1D). The reason for doing so was twofold: to see if both aging and young animals could utilize PPI as a detection metric, and to see if there were temporal deficits in sound processing abilities. There was significant effect of gap length on PPI value (F=9.272) (Figure 1E), but the differences in detectable gap length was similar between young and old gerbils (Figure 7F; young 15 ms: p=0.0103, 20 ms: p=0.0006; old gerbils 10 ms: p=0.0010, 15 ms: p=0.0004, 20 ms: p=0.0140). There was no significant effect of age on PPI value (F=3.062) or interaction between gap length and age on PPI value (F=0.6436), with the exception of the 20 ms gaps where we found that aging gerbils had a slight reduction of PPI values, indicating less detection of these 20 ms gaps compared to young gerbils (p=0.0365). These findings indicate that both young and aging gerbils can utilize PPI. Moreover, there may be some temporal impairment of gap detection at longer gap presentations.

Finally, we presented a broadband speaker swap of varying angles relative to midline to act as a prepulse prior to the startle stimuli (Figure 1F) to test spatial hearing abilities of young and aging gerbils. We presented 70 dB broadband noise at a speaker on one side of the animal’s midline and swapped this broadband noise to a corresponding identical speaker on the opposite side of the animal 80ms prior to the startle stimulus. The rational of this test was that animals would detect a speaker swap of wider angles more easily thus the swap angle would function as an assay of the animal’s binaural hearing abilities. Both angle (F=13.99) and age group (F=13.51) were sources of variation on PPI value. We found there was an interaction between age group and angle on PPI value (F=2.415) (Figure 1G). Specifically, young animals were able to detect speaker swaps of 30 degrees (p=0.0005) or larger (45 deg: p<0.0001, 90 deg: p<0.0001). By contrast, aging animals were only able to detect gaps of 45 (p=0.0043) and 90 degrees (p=0.0084). Additionally, we also measured reduced PPI values in aged animals compared to young animals at all angles (30 degrees: p=0.0041; 45 degrees: p=0.0105; and 90 degrees: p=0.0015). Smaller angle swaps were not well detected by either young or old gerbils, indicating that the detection of these narrow angle speaker swaps is challenging for both age groups. However, for wider angel speaker swaps, aging animals performed significantly worse than young animals. These results suggest that Mongolian gerbils recapitulate the age-related hearing deficits experienced by human listeners. In the next sections we will explore the mechanisms underlying these deficits in gerbils, because this animal model will allows us mechanistic and invasive studies which would not be possible in human listeners.

### Auditory brain stem responses are altered in old gerbils

To test the physiological responses of the auditory brain stem to sound, we performed auditory brain stem response (ABR) recordings, a non- or minimally invasive method that measures the compound activity of neurons in lower auditory centers in response to brief sound stimuli such as clicks. The rationale is that these clicks trigger action potentials in all auditory nerve fibers (spiral ganglion neurons) at nearly the same time, which results in a large compound event that can be measured from outside the skull as the first wave (peak) of the ABR trace. As this synchronized activity ascends the auditory pathway, it creates a series of additional waves that represent activity corresponding to different auditory brainstem nuclei. In gerbils, wave I is mediated by the spiral ganglion (auditory nerve) neurons, wave II is generated by of neurons in the cochlear nucleus, wave III is largely mediated by the medial nucleus of the trapezoid body (MNTB), wave IV corresponds to the output of the superior olivary complex, which contains the binaural lateral and medial superior olives, and wave V is created by the lateral lemniscus and inferior colliculus^49,50^.

All five ABR waves were distinctly present in most young (Figure 2A) and old gerbils (Figure 2B) when click stimuli were used. There was no significant difference in click ABR thresholds between young (N = 62) and old gerbils (N = 30). Thresholds were determined in two ways: First, by using automated algorithms to determine ABR thresholds (Young: 32.01 ± 12.71 dB SPL (mean ± SD); old: 29.86 ± 12.00 dB SPL; U = 1014.5, P = 0.4841 for a two-sided Mann– Whitney U test) (Figure 2C); or second, by manual inspection by an observer blinded to the experimental condition (Young: 31.13 ± 13.92 dB SPL; old: 34.00 ± 14.28 dB SPL; U = 872.5, P = 0.6271 for a two-sided Mann–Whitney U test) (Figure 2D). The probability that a particular wave was present in the ABR trace increased with increasing click sound intensity in both groups (Figure 2E and 2F). However, the young gerbils showed a somewhat higher probability of occurrence of wave I at high sound levels (young: 83.87% at 90 dB SPL; old: 73.33% at 90 dB SPL) and wave III at a medium sound level (young: 82.26% at 60 dB SPL; old: 66.67% at 60 dB SPL). These data imply that aging may affect cochlear and potentially MNTB activity to some extent, but the impact of increased hearing thresholds due to age-related hair cell loss in the inner ear was relatively minor.

**Figure 2.**
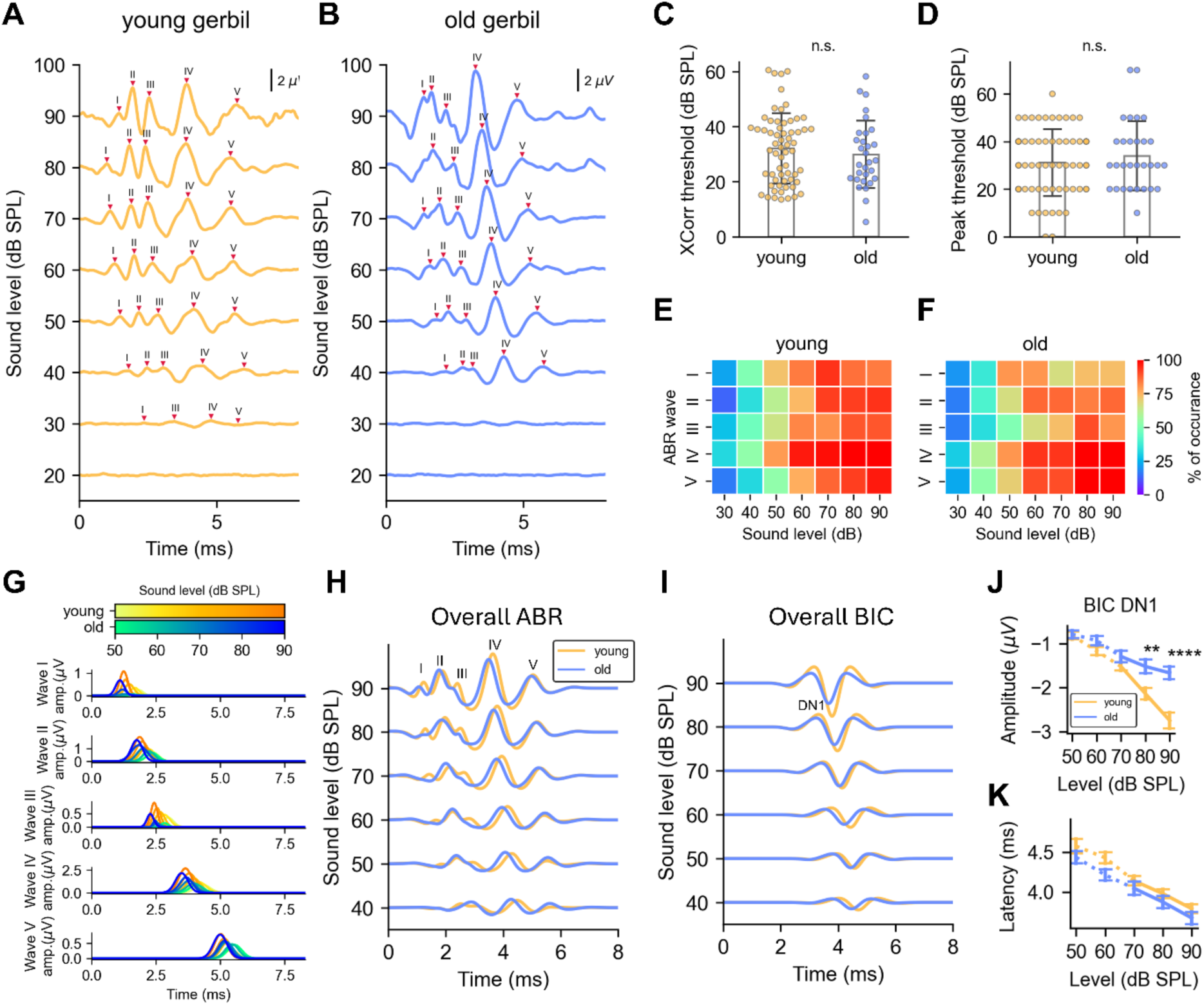
Click ABR and BIC traces in young and old gerbils. **(A)** Sample ABR traces recorded from a young animal (P85) elicited by click stimuli at different intensity levels. **(B)** Sample ABR traces from an old gerbil (P757). **(C)** ABR click thresholds calculated using the sigmoid function fitting on signal cross-correlations. (Young: 32.01 ± 12.71 dB SPL (mean ± SD); old: 29.86 ± 12.00 dB SPL (mean ± SD); U = 1014.5, P = 0.4841 for a two-sided Mann–Whitney U test). **(D)** ABR thresholds derived from a semi-automatic peak detection method used by blinded observers (Young: 31.13 ± 13.92 dB SPL (mean ± SD); old: 34.00 ± 14.28 dB SPL (mean ± SD); U = 872.5, P = 0.6271 for a two-sided Mann–Whitney U test). **(E-F)** Probability of occurrence of a particular ABR wave at a particular sound intensity when click sounds were used with young **(E)** and old gerbils **(F)**. **(G)** Overall ABR peaks (across animal grand mean) based on averaged peak amplitudes, widths and latencies in young (orange) and old (blue) gerbils. **(H-I)** Overall ABR **(H)** and BIC **(I)** traces (across animal grand mean) traces based on average peak metrics from young (orange) and old gerbils (blue).**(J-K)** Peak amplitudes **(J)** and peak latencies **(K)** of BIC-DN1 peak between young (N = 62; orange) and old (N = 30; blue) gerbils using Mann-Whitey U test (*: p<0.05; **: p <0.01; ***: p<0.001; ****: p<0.0001).

Next, we built an “overall ABR” which was based on mean metrics of the detected peaks across all animals. During this process we initially built single overall peaks (Figure 2G) and then entire mean overall ABR waveforms (Figure 2H). These waves were then used to explore differences between the young and the old age groups in more detail. Wave I and wave III were largely attenuated, and wave IV was advanced in old gerbils, but other waves were mostly consistent between the across-animal average waves of young and old animals. Despite the differences in wave I amplitude, the peak metrics of wave II remain unchanged, which could be caused by latent neural plasticity in the cochlear nucleus that compensates for hearing loss. Additional age-related changes were observed in the form of degraded wave III responses, suggesting degraded responses of MNTB activity, since MNTB is the main contributor to wave III in rodents.

Binaural processing was assessed by evaluating binaural interaction components (BICs)^47^. These were calculated for each recording through a point-by-point subtraction of the sum of the left plus right monaural ABRs minus the binaural ABR. If the lower auditory system (hypothetically) contained no binaural pathways, the sum of the monaural ABRs should be the same as the binaural ABR, resulting in a zero amplitude BIC trace. Any peaks in this trace, by contrast, indicate binaural processing ^51^. The prominent negative peak which can be seen in most BIC traces is the DN1 component (Figure 2I) and was quantified using the same method as for the ABR waves. The amplitude of the BIC-DN1 was significantly reduced in aging gerbils (N = 30) compared to young animals (N = 62) (Mann Whiteny U test; p < 0.01 at 80 dB; p < 0.0001 at 90 dB) (Figure 2J), with no significant difference in the BIC-DN1 peak latency (Figure 2K). This change in the BIC-DN1 amplitude suggests reduced binaural processing in old gerbils.

Several characteristic metrics of the five ABR waves are plotted in Figure 3. On average, old gerbils exhibited significantly attenuated wave I and wave III amplitudes (Figure 3A), indicating less synchronous firing and/or fewer contributing auditory nerve fibers and MNTB neurons to the responses. Interestingly, wave II did not show equivalent weaker responses in the older animals, suggesting that any age-related decreases in wave I (auditory nerve) have been compensated by circuits in the cochlear nucleus (wave II). If so, the age-related decreases in wave III reported here would have been created de novo in MNTB. Regarding peak latency, most waves showed only minor differences between young and old gerbils across all sound levels (Figure 3B). Further analysis on the peak intervals between the various ABR waves revealed no significant differences of peak latencies relative to wave I (Figure 3C). However, we found significantly shortened wave II-III and II-IV intervals in old gerbils at all sound levels (Figure 3D), suggesting a degradation of inhibitory activity especially between cochlear nucleus and superior olive.

**Figure 3.**
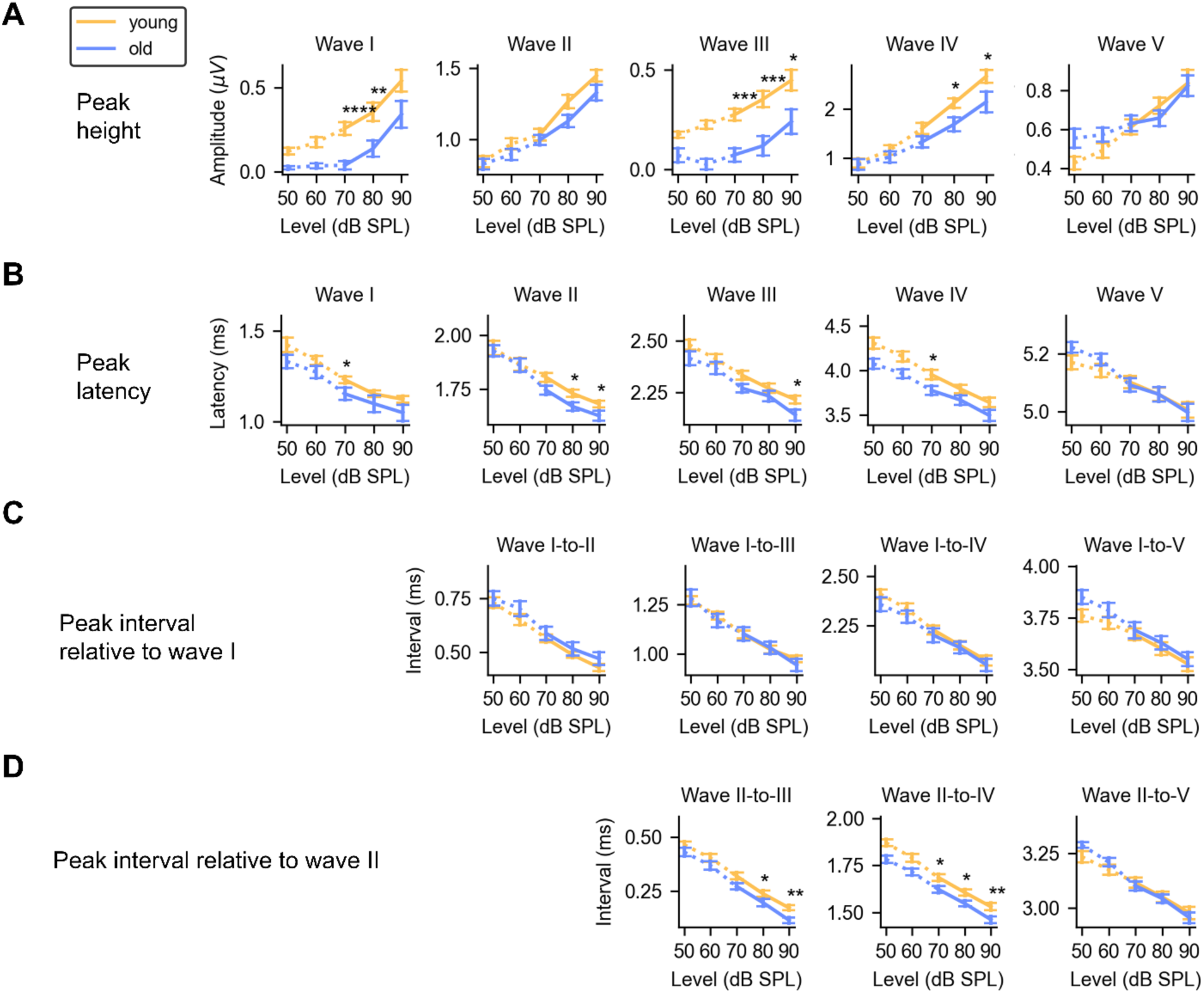
Peak metrics of ABR waves in young (N=62; orange) and old (N=30; blue) gerbils using the Mann-Whitey U test (*: p<0.05; **: p <0.01; ***: p<0.001; ****: p<0.0001). **(A)** Peak height. **(B)** Peak latency. **(C)** Peak interval relative to wave I. **(D)** Peak interval relative to wave II.

### CARS imaging shows decreased axon diameter in aging

After observing the electrophysiological abnormalities in the aging gerbil auditory brainstem, we hypothesized that these abnormalities could be attributed to changes in axonal conductance because previous work has shown the potential for plasticity of myelination throughout life^52–54^. Using CARS microscopy^40,41^, we first measured the axonal diameter of globular bushy cell axons in the auditory brainstem that innervate the MNTB and form the calyx of Held (Figure S1). In both young and old gerbils, the MNTB was sampled with multiple z-stack images, which were subsequently analyzed for axon thickness (Figure 4). We found a significant difference in axon width between the two age groups: the mean (±SD) axon diameter in young gerbils measured 3.9µm ± 0.18 (n=10), whereas in old gerbils it was reduced to 2.9µm ± 0.18 (n=7, p<0.0001, t-test, Figure 4C).

**Figure 4.**
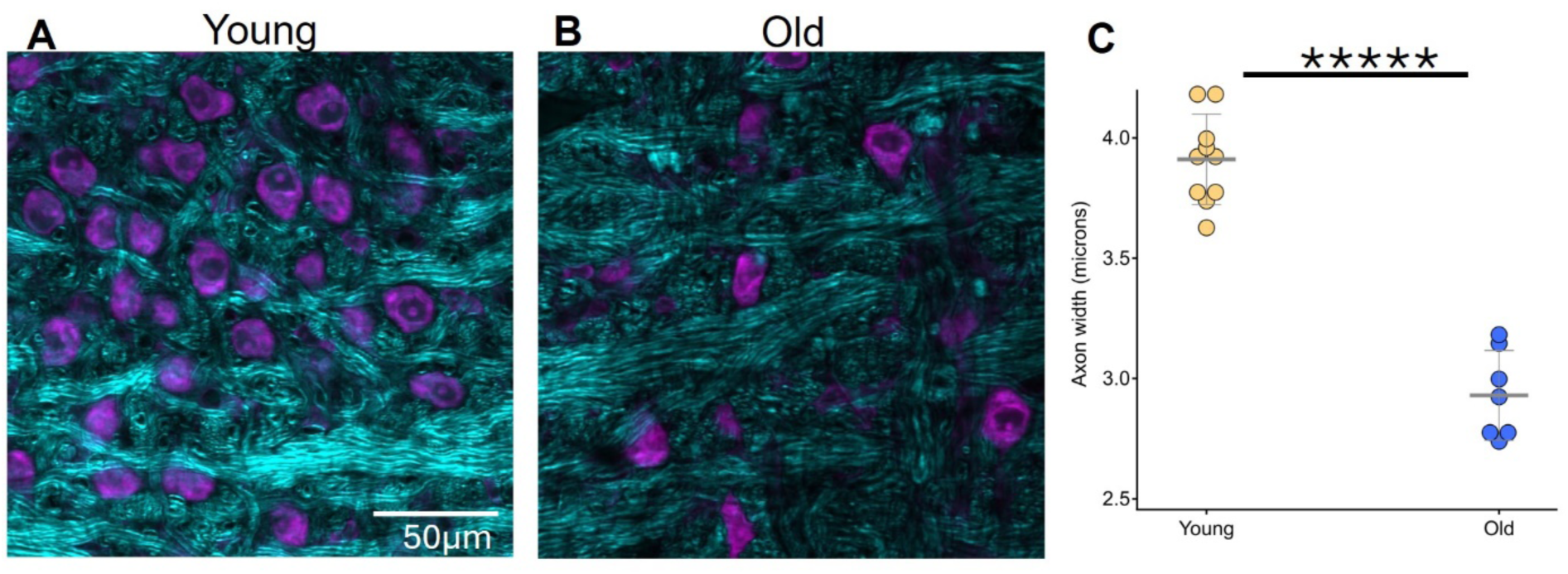
Fiber diameter size is reduced in old gerbils. Representative MNTB images show cell bodies stained with Nissl (purple) and CARS signal (cyan) in young **(A)** and old **(B)** gerbils. **(C)** Box plots represent data from individual animals (n=10 young, n=7 old, with 30 measurements per section and 2–6 sections per animal). ****p = 7e-8

### EM suggests decreased myelination in old animals, altering conduction velocity

A total of 3027 myelinated axons were sampled and quantified in 133 EM images captured from 6 gerbils (Young: n_axon_ = 1266, N_animal_ = 4; Old: n_axon_ = 1761; N_animal_ = 2) (Figure S1). The two groups presented several differences in the cross-section of axon fibers (Figure 5A), with young gerbils containing thicker axon fibers with clearer and better-defined myelin layers. Age related changes were also evident in CARS images, in particular a reduction in fiber diameter (Figure 5D; young: 2.948 ± 1.22 µm (mean ± SD), old: 2.349 ± 1.271 µm; P < 0.0001, Mann–Whitney U test) and myelin thickness (Figure 5E; young: 0.608 ± 0.242; old: 0.561 ± 0.304 µm; P < 0.0001, Mann– Whitney U test). Thus, the average fiber diameter was significantly decreased by about 0.6 µm, and the myelin thickness decreased by around 0.05 µm.

**Figure 5.**
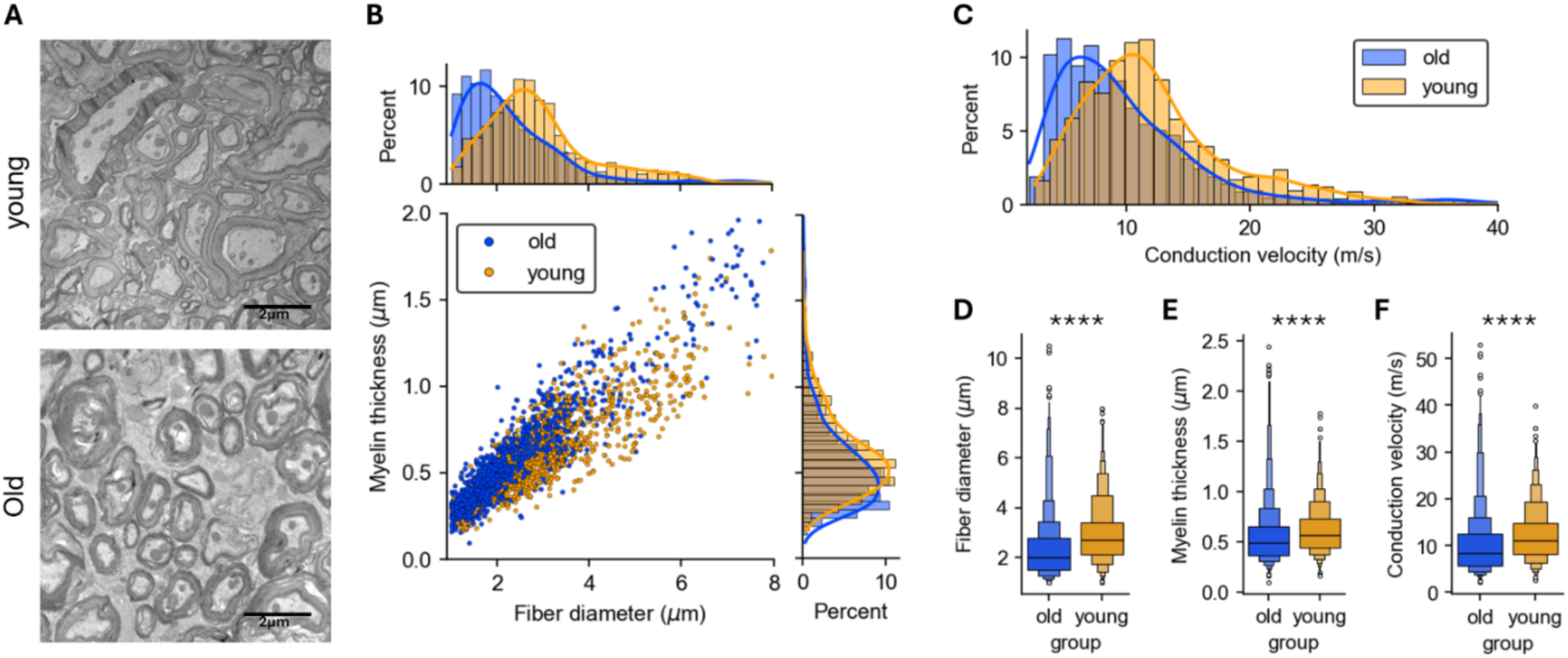
Myelination properties acquired from EM images. **(A)** Example EM images captured from a young gerbil (top) and an old gerbil (bottom). (error bar: 2 µm) **(B)** Fiber diameter and myelin thickness of quantified axon fibers sampled from the young group (n_axon_ = 1266; N_animal_ = 4) and the old group (n_axon_ = 1761; N_animal_ = 2). **(C)** Distribution of the fiber conduction velocity estimated by simulating the action potential propagation along myelinated axons. **(D)** Fiber diameters in two groups (Young: 2.948 ± 1.22 µm (mean ± SD), 2.707 µm (median); Old: 2.349 ± 1.271 µm (mean ± SD), 2.022 µm (median); P < 0.0001 in the two-sided Mann–Whitney U test). **(E)** Myelin thickness in two groups (Young: 0.608 ± 0.242 µm (mean ± SD), 0.563 µm (median); Old: 0.561 ± 0.304 µm (mean ± SD), 0.49 µm (median); P < 0.0001 in the two-sided Mann– Whitney U test). **(F)** Estimated fiber conduction velocity in two groups (Young: 12.224 ± 5.90 m/s (mean ± SD), 11.143 m/s (median); Old: 10.145 ± 6.783 m/s (mean ± SD), 8.435 m/s (median); P < 0.0001 in the two-sided Mann–Whitney U test).

We modeled the physiological effects of this reduced myelin thickness with a myelinated axon model that has been used previously for this neural circuit^18,55^ and found a slower conduction velocity of action potential propagation along axons of older animals (Figure 5F; young: 12.224 ± 5.90 m/s; old: 10.145 ± 6.783 m/s; P < 0.0001, Mann–Whitney U test), with the fiber conduction velocity slowed by 1.081 m/s. However, these metrics displayed skewed distributions (Figure 5B and 5C), and the centroids of the histograms were clearly separated with median differences of 0.685 µm in fiber diameter and 0.073 µm in myelin thickness, resulting in 2.703 m/s in conduction velocity per the models. Note that the larger variations in the EM fiber metrics of the old animals (Figure 5D-5F) translated with our model into an increased standard deviation of the conduction velocity, suggesting a relatively larger variation of transmission latencies in the old group (± 204 µs for 3-mm distance) compared to the young group (± 177 µs for 3-mm distance). Consequently, the axon fibers of old gerbils show degraded action potential propagation with longer and more variable transmission latencies. By calculating the time course of action potential propagation through a 3-mm axon fiber with conduction velocities between the 25% to 75% quartile levels, we calculated that approximated transmission latencies range from 0.20 to 0.37 ms in young gerbils and from 0.24 to 0.52 ms in old gerbils.

### Oligodendrocyte loss in aging contributes to the myelin deficits

Using EM and CARS microscopy, we identified an age-dependent decrease in myelination. Oligodendrocytes (OLs) are specialized glia cells responsible for producing myelin which insulates axons in the central nervous system, and the loss of these cells can lead to changes in myelination of MNTB afferents^41^. To investigate whether the age-related myelin changes are linked to alterations in the quantity of mature (OL) and precursor oligodendrocytes (OPC), we employed an immunohistochemical method. Representative z-stacks of aspartoacylase (ASPA) and sex determining region Y-box 10 (SOX-10) labeled MNTB tissue were imaged with a confocal microscope. SOX-10 serves as a marker for the entire OL lineage (OPC + OL), whereas ASPA specifically identifies mature OLs. Thus, cells solely stained for SOX-10 represent OPCs, while co-staining with ASPA indicates mature OLs (Figure 6). Our initial focus was on comparing the density and maturation status of MNTB OLs across different age groups. The rationale of this experiment was that only mature OLs produce myelin but not immature or precursor cells and therefore, a decreased count of mature oligodendrocytes in aging might contribute to the reduced myelination observed in the older age group.

**Figure 6.**
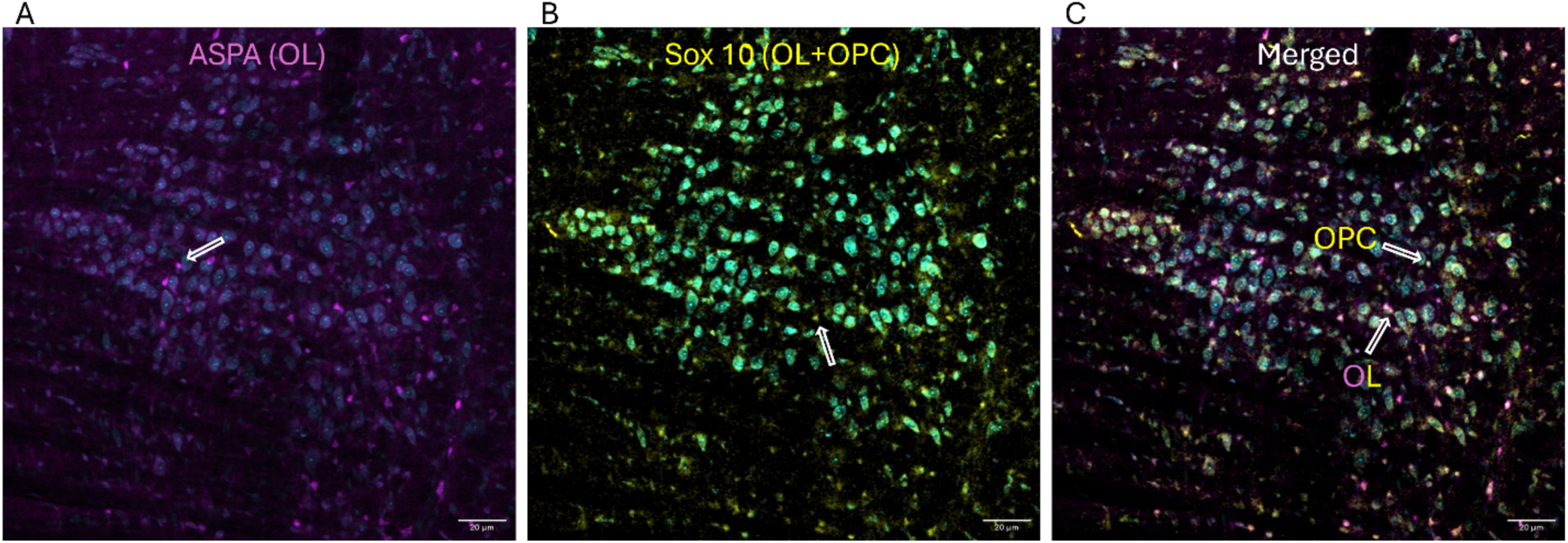
Oligodendrocytes (OLs) and Oligodendrocyte Precursor Cells (OPCs); **(A)** ASPA, a specific marker for mature OLs, is shown in magenta. **(B)** Sox-10, a marker for the entire OL lineage (OPCs and OLs), is labeled in yellow. **(C)** OPCs were measured from cells labeled with Sox-10 but not ASPA. Mature OLs express both markers. MNTB cell bodies are indicated in cyan (Nissl).

We found a significant and substantial reduction in ASPA signal within the aging population (1595 ± 91 vs 473 ± 171, p=0.0005, for young and aged gerbils, respectively, Figure 7C). Interestingly, the total number of oligodendrocyte precursor cells remained relatively consistent (Figure 7D), suggesting the potential for the regeneration of lost oligodendrocytes from parenchymal OPCs. These findings suggest an age-related loss of oligodendrocytes results in myelination deficits.

**Figure 7.**
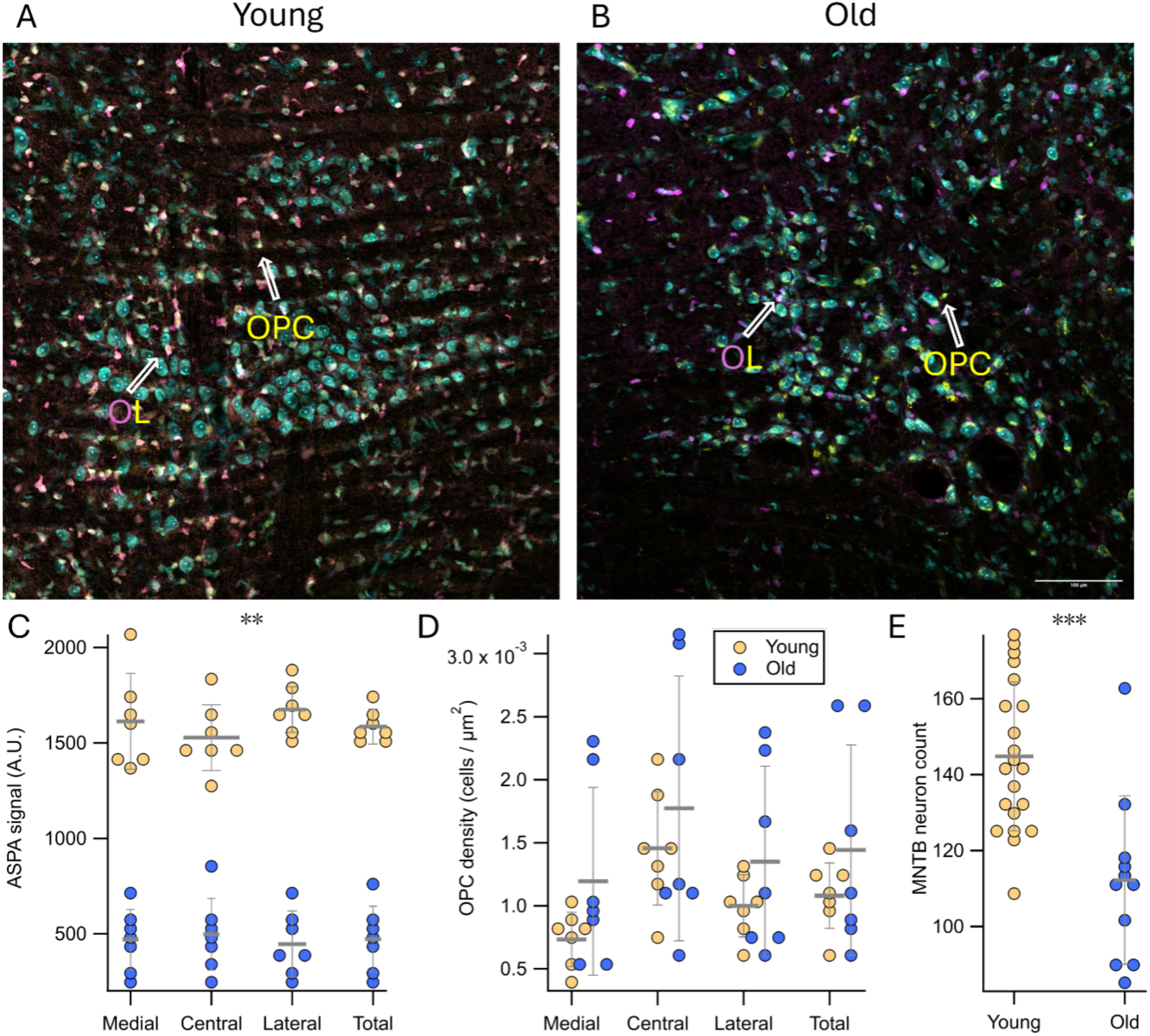
Oligodendrocyte (OL) numbers decrease compared to oligodendrocyte precursor cells (OPC), in the MNTB of old gerbils. Representative images show MNTB cells (cyan), OLs (ASPA/SOX-10 marker in magenta/yellow) and OPCs (SOX-10 without ASPA staining) in young **(A)** and old **(B)** gerbils. **(C)** ASPA fluorescence levels for young (yellow, n=6) and old (blue, n=6) gerbils across total, medial, center, and lateral MNTB. ** p<0.01 using two-tailed Mann-Whitney test and Bonferroni correction for multiple comparisons. No difference was found across MNTB tonotopic areas within each age group. **(D)** OL density in MNTB is similar in both age groups. **(E)** Estimated number of MNTB neurons (teal cells in A and B) per 640 µm * 640 µm region. Error bars: standard deviation. *** p<0.001 using two-tailed Mann-Whitney test.

The MNTB tonotopic axis encodes high to low sound frequencies across the medial to lateral dimension. Previous research found that parameters such as soma size, synaptic properties and myelination change along this tonotopic axis^18,20,56^. Accordingly, we assessed OL in three MNTB divisions: medial (high frequency), central (intermediate frequency) and lateral (low frequency)^41^. We found a substantial reduction in OL numbers between young and old gerbils within each tonotopic section (Figure 7C). However, we found no significant changes in either combined OL + OPC counts or the counts of each individual cell type between the tonotopic regions within the same age (Figure 7C and 7D).

Next, we performed cell counts of MNTB neurons to determine whether the age-related reduction of mature oligodendrocytes could be explained by an age-related reduction in neurons in the same brain nucleus. The rationale was that fewer neurons might require fewer oligodendrocytes to myelinate the axons of these neurons. We did find, in fact, a slight age-related reduction in the number of MNTB neurons (Old: 110.17 ± 25.62 cells, N_animal_ = 11; Young: 143.47 ± 25.62 cells, N_animal_ = 21; p = 0.0006, Mann-Whiteney U test; Figure 7E). However, this ∼20% reduction in neurons was much smaller than the observed ∼300% reduction in mature oligodendrocytes, suggesting that the ratio between neurons and mature Ols was decreased in the older age group.

## DISCUSSION

We report five main findings: First, physiological responses of the auditory brain stem, specifically nuclei of the auditory brainstem responsible for the initial encoding of the acoustical cues to sound location, change with age. The largest age-related changes we observed in ABR recordings were in waves I, III and the BIC of the ABR. Wave III largely reflect neural activity in the MNTB^57–59^, thus the changes in this wave indicate age-related changes with MNTB physiology. Importantly, while we also found age-related changes in wave I, the observed changes in wave III seem to be independent rather than a simple reflection of the wave I changes. These results are wholly consistent with prior age-related studies in gerbil ABR waves^60^. Second, the exceptionally heavily myelinated afferent axons to MNTB show age-related decreases in myelination, both with CARS and with EM imaging. Third, when the physiological consequences of the observed decreases in myelination were modeled, the modeling results were consistent with the empirical data obtained from ABR recordings, indicating that reduced myelination is a main factor in this altered physiology. Fourth, the number of mature oligodendrocytes was reduced in old gerbils, consistent with the reduction in myelination. However, the *total* number of oligodendrocyte precursor cells was unchanged, suggesting an age-related impairment in oligodendrocyte regeneration. Fifth, this demyelination of MNTB afferent fibers has consequences for the sound localization and spatial hearing abilities of affected animals, which was measured as impaired detection of spatial cues in a behavioral task.

We suggest that the wave I reductions cannot explain the wave III reductions, because wave II was unchanged between young and old animals. In other words, any peripheral hearing loss that the older animals may have experienced was not measurable at the level of the cochlear nucleus. A potential reason for this is the presence of compensatory mechanisms which upregulate auditory gain. Such mechanisms are commonly observed in many forms of hearing loss and might explain why wave II amplitudes were similar between young and old animals^61^. If that is the case, the wave III reductions observed in old animals must have a mechanism which is at least partially independent of peripheral hearing loss. Alternatively, it is possible that the CN neurons that generate wave II are a different population than the globular bushy cells that provide the input to MNTB and thus wave III. We show that the globular bushy cell axons to MNTB are significantly demyelinated. As a consequence of this, there is more jitter in the input, thus causing a desynchronized response from MNTB leading to reduced wave III (and thus reduced BIC). Because of the giant calyx, the only strong reason for reduced wave III is because the inputs to MNTB are also highly jittered due to demyelination.

The mammalian sound localization pathway critically depends on the very accurate timing of the arrival of sound information at auditory brainstem nuclei^16^. Sound localization is a computational process which relies on the comparison of arrival time and intensity of the same sound wave between the two ears. Low frequency sounds are localized by the interaural time difference, which can be as small as only several tens of microseconds at the system’s localization threshold, about two orders of magnitude faster than a typical action potential^16,62^. Not surprisingly, there are many specializations for the processing of neural information with very high levels of temporal fidelity. Perhaps the most extreme examples of these specializations can be found in the pathway of the MNTB. Principal neurons in MNTB feature several adaptations to process information with very high levels of temporal fidelity. One such adaptation is the calyx of Held^63,64^, a type of giant synapse which innervates the MNTB principal cell body. Furthermore, MNTB neurons have small dendrites, are electrically very compact, have large synaptic currents, a low input resistance, and kinetically fast receptors, just to name a few of these subcellular specializations^17,65,66^. The most likely functional significance of all these specializations is speed and temporal fidelity in MNTB activity, which is obviously required by the neural targets of MNTB. The target nuclei of MNTB include but are not limited to the principal sound localization nuclei, the lateral and medial superior olive (LSO and MSO respectively)^67^. Since MNTB neurons are glycinergic ^68,69^, the role of MNTB within the sound localization process is to provide fast and well-timed neural inhibition to the sound localization process, which itself is critically dependent on temporal precision.

Consistent with the reduction in wave III ABR amplitudes, we also found that the BIC of the ABR was significantly reduced indicating compromised binaural coding of ITD cues. Recent evidence suggests that the BIC is produced by the LSO^51,70,71^. The BIC has been used as a biomarker for binaural hearing performance and reduced or eliminated BIC has been correlated with poor binaural hearing performance in humans in a variety of different hearing pathologies, including aging^47^.

In this context it seems intuitive that afferent axons to MNTB also need to be capable of transmitting neural activity quickly and with high temporal fidelity. These axons are therefore exceptionally thickly myelinated and additionally, the spacing and size of the nodes and internodes is tightly controlled to accomplish particular action potential propagation time constants^18,20^. Moreover, sound elicited neural activity in this pathway has been shown to contribute to the establishment of these tight parameters in an activity dependent way during postnatal development^27^.

Our results indicate that this axon population partially demyelinates during aging. Equally important, the population of axons in older animals has a wider distribution of myelination levels, i.e. some axons are more demyelinated than others. Reduced myelin thickness should lead to reduced capabilities to conduct action potentials as well as increased jitter in these fibers, causing the high level of temporal fidelity in these afferents to be lost. Additionally, normal aging has been associated with a partial loss of myelin in other neural systems, and remyelination can result in shorter internode length and decreased sheath thickness, which would also decrease the action potential propagation speed. It would additionally compromise the extreme temporal fidelity and precision that was established in this system during early postnatal development, and that is required for accurate sound localization^72^. Our model-estimated conduction velocity based on myelination data from EM images further supports this idea by suggesting an increased variability of the fiber conduction velocity and prolonged transmission latencies in older animals.

What might be the mechanisms for this age-related demyelination of the MNTB afferent fiber bundle? One possibility is that general age-related myelin degeneration and incomplete regeneration may play a role here, as has been demonstrated in other neural systems^72,73^. Our results also show a decrease in mature oligodendrocytes in aged animals, which would be consistent with incomplete oligodendrocyte regeneration^74^. Interestingly, while we observed an age-related decrease in mature oligodendrocytes, the total number of oligodendrocyte precursors was comparable to young animals, suggesting that regeneration of new oligodendrocytes during the animal’s lifetime might be an important mechanism of myelin degeneration with age.

Another well-described age-related change in the peripheral auditory system is synaptopathy – the progressive loss of hair cell to auditory nerve fiber synapses with age. A physiological indicator of cochlear synaptopathy in rodents is a reduction in the amplitude of ABR wave I relative to controls^75^. Our finding of reduced amplitude wave I in aged gerbils is wholly consistent with these animals having synaptopathy despite normal hearing thresholds. While this loss may not necessarily lead to increased auditory thresholds, it could plausibly lead to decreased activity levels in downstream auditory areas such as the cochlear nucleus and MNTB^76,77^. Potentially, this chronically reduced activity may contribute to reduced myelination. One mechanism for reduced activity in brainstem neurons might be caused by the loss of subthreshold inputs to cochlear nucleus bushy cells from auditory nerve due to synaptopathy. It is well known that convergence of auditory nerve fibers onto cochlear nucleus bushy cells is important for the emergence of precise coding of both temporal fine structure and envelope cues in the brainstem for suprathreshold sounds^78^. Computational modeling by Budak et al.^79^ demonstrates that synaptopathy comparable to that found empirically in animal models leads to significant lowering of overall spike rate in globular (GBC) and spherical (SBC) bushy cells of the cochlear nucleus as well as the medial superior olive (MSO) by anywhere from 30-70%. SBCs from both ears provide excitatory inputs to the MSO, which encodes low-frequency fine structure interaural time differences (ITDs). SBCs provide the excitatory input to the lateral superior olive (LSO) from the ipsilateral ear while GBCs from the contralateral ear provide inhibitory input via a synapse through the MNTB. Budak et al.^79^ also showed that synaptopathy significantly reduced phase locking in SBCs and GBCs and thus the ability of the simulated MSO to encode the ITD cue was severely degraded. Our finding here of age-related reductions in myelin in brainstem axons despite no significant hearing loss is consistent with a hypothesis that activity-dependent reductions in spiking of cochlear nucleus bushy cells as well as phase locking due to synaptopathy^79^ could plausibly lead to reduced myelin over time. Reduced myelin would be expected to lead to increased spike jitter and thus poorer temporal encoding of sound, which in turn would cause substantial deficits in the encoding of the acoustical cues to sound location by neurons in the MSO and LSO. Indeed, our demonstration here of reduced amplitude of the BIC shows that the jitter caused by demyelination to the inputs to MNTB can cause impaired binaural cue coding by LSO. This, then, would be expected to cause difficulties in binaural and spatial hearing behavioral tasks, such as sound localization or listening in noisy environments.

Consistent with this view are recent results from oligodendrocytes in the developing auditory system, which show that one population of these cells contains voltage gated sodium channels, is excitable, and that excitation of this population supports maturation of these cells^80^. While a parallel role in aging has not been shown, it seems possible that reduced activity from the auditory nerve may further contribute to the impaired oligodendrocyte regeneration and, by extension, the impaired myelination of axons.

We note that there are some striking similarities between this body of work and recent work from the autism field. Autistic human listeners, as well as animal models of autism (the Fragile X mouse model, representing the most common monogenetic cause for autism), also have reduced ABR wave III^81,82^. Moreover, MNTB afferent axons from Fragile X mice show a similar reduction of myelination as we show here for aging^41^. Additionally, the same Fragile X mice also have an increased population of mature oligodendrocytes and precursors^41^. While synaptophathy and hidden hearing loss may play a role in the aging population, it is less likely this mechanism would also be active in the Fragile X mice, which are all young adults. In Fragile X, the underlying mechanism would more likely be impaired development, leading to aberrant oligodendrocyte maturation, resulting in incorrect myelination of the axon bundle. Additionally, Fragile X mice showed similar impairment in spatial hearing as measured by PPI ^83^, similar to what was seen in our aging gerbils. The overall outcome is similar, since both autistic and aging listeners experience similar sound localization deficits and cocktail party performance deficits, while also having the same compromised temporal fidelity in MNTB afferent pathways.

## ACKNOWLEDGEMENTS

We would like to thank Dr. Anza Darehsouri for help with electron microscopy, Mr. Carol Mirita and Dr. Dominik Stich for help with CARS microscopy, and Dr. Tim Lei for designing and writing the software used to measure auditory brain stem responses. We would also like to thank Dr. Alon Poleg-Polsky for many helpful discussions and for support with the oligodendrocyte imaging and quantification.

## AUTHOR CONTRIBUTIONS

S.P., B.-Z.L., M.S., M.R. designed study, collected data, analyzed data, performed modeling. E.G.H., D.J.T. and A.K. conceived of study, designed experiments.

## METHODS

All animal procedures conducted in this study were approved by the Institutional Animal Care and Use Committee (IACUC) of the University of Colorado Anschutz Medical Campus, permit number 00617, and followed all OLAW and National Institutes of Health (NIH) standards for the humane treatment of laboratory animals. A total of 92 gerbils were used in this study. The animals were raised and aged in-house. Animals from both sexes were used in the experiments for young (76 ± 17 days) and old (2 years 7 month, 930 days ± 3 month).

### Auditory brainstem response (ABR) acquisition

ABR recordings were performed following procedures similar to those detailed in previous studies^83,84^. Briefly, gerbils were anesthetized using a combination of 100 mg/kg ketamine and 10 mg/kg xylazine, then placed on a heating pad inside a small soundproof chamber. The stimulus presentation, generation, and recording of evoked potentials were managed through custom Python software interfacing with an RME Fireface UCX Sound Card (REM audio, Haimhausen, Germany) operating at a sampling rate of 44.1 kHz. Stimuli were delivered via calibrated Etymotic ER10B ear coupling tubes and ER2 earphones (Etymotic, Elk Grove Village, IL, USA) using standard sound delivery tubes. Evoked potentials were recorded using platinum subdermal needle electrodes simultaneously placed at the apex and behind each pinna (active), referenced to the nape of the neck, grounded with a hind leg ground channel. The signals were then amplified, digitized, and processed for output using a range of specialized equipment including a Tucker Davis Technologies (TDT) low-impedance head stage (RA4LI), preamplifier (RA4PA), and digital amplifier (x10,000), along with a multi-input/output processor (RZ5) (Tucker-Davis Technologies, Alachua, FL, USA).

ABRs were elicited in response to 500 repetitions of sound stimuli consisting of 0.2 ms monaural (left or right) and binaural complex clicks with interstimulus intervals of 35 ± 5ms (mean ± SD). The responses were initially recorded for 90 dB SPL stimulus presentations and then gradually decreased in 10 dB SPL steps until the threshold level was reached. The threshold was defined as the average of the minimally detectable waveform and the next dB SPL step where a discernible ABR signal emerged in any of the three recorded channels.

### ABR analysis

The recorded raw ABR traces were first band-pass filtered with a 100-3000 Hz 2nd order Butterworth filter and then smoothed and detrended using a Savitzky-Golay filter. Wave peaks were automatically detected and labeled by finding the local maxima on the processed ABR traces, and the wave labels were then inspected and adjusted if misclassified. All labeled peaks were quantified by the peak height, the peak half-width, and the peak latency.

Click ABR thresholds were estimated using two different approaches. For the objective estimation method, the cross-correlation (xCorr) of ABR traces between adjacent sound levels was calculated. The xCorr thresholds were computed by fitting the correlation-coefficient vs. sound level data to the sigmoid function^85^. For the conventional estimation method, the peak thresholds were determined by the sound level at 10 dB SPL below the minimum sound level that elicited visually detectable wave peaks. Researchers analyzing the ABR data were blinded to the animal and group information.

### Tissue Preparation

Gerbils were euthanized via an overdose of pentobarbital (120 mg/kg body weight Fatal+, Vortech Pharmeceuticals, Dearborn, MI, USA and subsequently transcardially perfused with phosphate-buffered saline (PBS; containing 137 mM NaCl, 2.7 mM KCl, 1.76 mM KH2PO4, 10 mM Na2HPO4, Sigma-Aldrich) followed by 4% paraformaldehyde (PFA). Following perfusion, the animals were decapitated, and their brains were removed from the skull. The extracted brains were then post-fixed in 4% PFA overnight before being transferred to 30% sucrose for further processing. After the brains sank, they were washed three times in PBS to remove residual sucrose. Brains were then put in a mold in OCT compound (SAKURA 4583, Sakura Finetek, Torrance, CA, USA) and left to freeze in a cryostat (Leica CM1950, Leica Biosystems, Nussloch, Germany) for 20 minutes. The brainstem was sliced to 50 microns for CARS imaging and IHC (Immunohistochemistry). Slices were collected starting before the 7th nerve.

For EM imaging, brainstems from six animals were embedded in 4% agarose (in PBS) immediately following perfusion. Brains were coronally sliced at 800 μm using a Vibratome (Leica VT1000s), with subsequent division through the midline. One half of the brainstem was processed for CARS and IHC; the other half was fixed in 2.5% glutaraldehyde, 2% PFA, and 0.1 M cacodylate buffer (pH 7.4) for 24 hours. The tissue was rinsed in cacodylate buffer (recipe) and immersed two times in a mixture of 1% osmium and 0.8% potassium ferrocyanide (Sigma Chemical), each for 1.5 hours. Following rinses in water, the tissue was stained with 4% uranyl acetate in 50% ethanol for 2 hours. The tissue was then dehydrated through a graded series of ethanol, embedded in Embed812 resin (Electron Microscopy Sciences, Hatfield, PA, USA), and cured for 24 hours at 60°C. Ultra-thin sections (65 nm) were cut with a diamond knife (Diatome, Quakertown, PA, USAon a UC7 ultramicrotome (Leica Microsystems, Wetzlar, Germany) and collected on 200 mesh copper grids for analysis. Images were acquired on a Tecnai T12 transmission electron microscope (ThermoFisher Scientific, Waltham, MA, USA) equipped with a LaB6 source at 120kV, using an NS15 (15 Mpix) camera (AMT Imaging, Woburn, MA, USA).

For myelination analysis with CARS (Coherent Anti-stokes Raman Spectroscopy) microscopy, two to five slices per brain were stained with Nissl (Neurotrace 425/435 Blue-Fluorescent Nissl Stain, Invitrogen, 1:200), in antibody media (AB media: 0.1 M phosphate buffer (PB: 50 mM KH2PO4, 150 mM Na2HPO4), 150 mM NaCl, 3 mM Triton-X, 1% bovine serum albumin (BSA)) for 30 minutes at room temperature on a standard laboratory shaker, then stored free-floating at 4°C until imaged.

### Immunohistochemistry

For staining oligodendrocytes, six to eight free-floating sections from each brain were submerged in L.A.B. solution (Polysciences, Warrington, PA, USA) for 10 minutes to expose epitopes. Sections were then washed twice in PBS and blocked in a solution containing 0.3% Triton-X, 5% normal goat serum (NGS), and PBS for 1 hour on a laboratory shaker. After blocking, sections were stained overnight with primary antibodies (Table S1): rabbit anti-Aspartoacylase (GeneTex, Irvine, CA, USA 1:1,000) and mouse Sox-10 (Santa Cruz Biotechnology, Santa Cruz, CA, USA, sc-365692, 1:500) in blocking solution with 1% NGS. The sections were then washed three times in PBS (10 minutes each wash) and incubated for 2 hours in secondary antibodies (Table S2) (diluted in blocking solution with 1% NGS). After three additional PBS washes (5 minutes each), sections were stained with Neurotrace 425/435 Blue-Fluorescent Nissl Stain (ThermoFisher Scientific, Waltham MA, USA, 1:200) in antibody media (0.1 M phosphate buffer, 150 mM NaCl, 3 mM Triton-X, 1% BSA) for 30 minutes. Finally, the stained sections were washed briefly in PBS, slide-mounted with Fluoromount-G (Southern Biotech, Birmingham, AL, USA), and stored at 4°C. All antibody labeling was performed at room temperature.

**Table S1.** Primary antibodies used in immunofluorescence.

| Antibody | Immunogen | Manufacturer, species, mono or polyclonal, cat. or lot no., RRID | Concentration |
| --- | --- | --- | --- |
| Aspartoacylase (ASPA) | Recombinant protein encompassing a sequence within the center region of human Aspartoacylase | GeneTex (Irvine, CA, United States), rabbit polyclonal, GTX113389, RRID:AB_10727411 | 1:1,000 |
| Sox-10 (A-2) | An epitope between amino acids 2-29 at the N-terminus of Sox-10 protein. | Santa Cruz Biotechnology, (Dallas, TX, United States), mouse monoclonal, sc-365692; RRID:AB_10844002 | 1:500 |

**Table S2.** Secondary antibodies used in immunofluorescence.

| Catalog No., RRID | Manufacturer | Reactivity | Conjugate | Concentration |
| --- | --- | --- | --- | --- |
| A-21428, AB_2535849 | Invitrogen | Goat anti-Rabbit IgG (H + L) | Alexa Fluor 555 | 1:500 |
| A-21235, AB_2535804 | Invitrogen | Goat anti-Mouse IgG (H + L) | Alexa Fluor 647 | 1:500 |

### Antibody Characterization

The primary antibody for mature oligodendrocytes, Aspartoacylase (ASPA, 1:1,000, GeneTex, Irvine, CA, USA; GTX113389), is a rabbit polyclonal antibody specific to mature oligodendrocytes (OLs). This antibody targets a recombinant protein sequence within the center region of human aspartoacylase, which converts N-acetyl-L-aspartic acid (NAA) to aspartate and acetate, aiding in white matter maintenance^86^. It has been validated for labeling mature OLs in mouse cerebral cortex through protein overexpression and western blot analysis^87,88^ as well as in our previous study of the auditory brainstem^41^ The Sox-10 antibody [Sox-10 (A-2), 1:500, Santa Cruz Biotechnology Santa Cruz, CA, USA, sc-365692] is a mouse monoclonal antibody specific to an epitope between amino acids 2–29 at the N-terminus of the human Sox-10 gene. This antibody labels the entire oligodendrocyte lineage (OPCs and OLs) and has been shown to specifically mark these cells in the CNS and mouse brains^89–91^. Both primary antibodies were visualized using fluorescent-conjugated secondary antibodies (listed in Table S2).

### Imaging

Brainstem slides for immunofluorescence were imaged using an Olympus FV1000 confocal microscope (Olympus) with 488, 543, and 635 nm lasers. The MNTB was identified by its trapezoidal morphology and cell size, and z-stacks were taken with a 20x objective (UPLSAPO20X, Olympus Life Science, Waltham, MA, USA NA 0.75) to visualize the entire nucleus. The nucleus was divided into medial, central, and lateral regions, following^56^. Briefly, the MNTB was digitally extracted using FIJI software, and the tonotopic axis was estimated by drawing a dorsomedial-to-ventrolateral line through the nucleus. This line was measured and divided into thirds with perpendicular lines to delineate the regions.

For CARS microscopy, brainstem sections were imaged using an Olympus FV-1000 with non-descanned detectors in both forward and epi CARS directions. Sections were placed in a culture dish with a coverslip bottom and held down by a custom glass weight. Z-stacks were taken with an Olympus 30x, 1.05 NA silicone oil objective (Olympus Life Science, Waltham, MA, USA) to collect the CARS signal in the epi direction. Signal in the forward direction was collected through an Olympus 0.55 NA condenser. An APE picoEmerald OPO laser system (APE, Berlin, Germany) was used to generate the pump/probe beam which is tunable from 770nm to 990nm, and the fixed wavelength Stokes beam at 1031 nm. Myelin was visualized by imaging the CH2 stretch-mode at 2,845 cm^−1^ and the pump/probe beam was set to 797.2 nm, resulting in a CARS signal at 649.8 nm. The tunable pump/probe beam also generates two-photon excitation fluorescence which was separated from the CARS signal by a dichroic mirror and detected in a separate channel by the non-descanned detector (NDD) unit in epi-direction.

Four brains from young and two from old gerbils were processed and imaged with electron microscopy. Sections for EM were first imaged on a compound microscope to identify the MNTB by cell size, shape, and location in parasagittal slices. Once located, slices were imaged on a FEI Tecnai G2 transmission electron microscope (ThermoFisher Scientific, Waltham, MA, USA) with an AMT digital camera (AMT Imaging, Woburn, MA, USA).

### Cell Counting

Quantification of OL’s and entire OL lineage (OPCs and OLs) was performed as follows. In FIJI, each stack was brightened, the background subtracted, the image scaled and the MNTB digitally extracted. We found nonspecific staining of MNTB cells by SOX-10 in aged animals; thus, to identify OL lineage cells we subtracted the Nissl image from the SOX-10 channel. Next, each slice of the SOX-10 channel was thresholded and the morphology of the (OL + OPC) cells within the MNTB was identified with FIJI particle analysis. Some cells were visible on neighboring slices. To prevent overcounting, we identified and combined close (less than 4 micron apart) particles into single cells with a custom-made script (Igor Pro). Masks were calculated for each cell, and the mean signal on the ASPA channel within the mask was recorded.

The number of MNTB neurons was automatically counted with a customized script using the scikit-image library^92^. For each 640 µm × 640 µm confocal image, the central z-stack was binarized, subtracted by the background obtained by the top-hat transform, and processed with a morphological erosion operation. The number of neurons of the image was approximated by counting the number of contours detected in the processed image. The MNTB neuron count for each animal was calculated as the average MNTB neuron count across all acquired images, with 4.09 ± 1.35 images (mean ± SD) acquired per animal.

### Myelination Analysis

Myelin diameter was measured from CARS images using FIJI software. The line tool was used to measure the fiber diameter for 30 axons per image (Figure S1A). Due to resolution limitations, CARS images could not be used to quantify myelin thickness or g-ratios.

For EM images, cross-sections of myelinated fibers were blindly sampled through visual inspection. The myelin layers were semi-automatically segmented by active contours. Dead axons with incomplete and loose lamellae as well as small fibers with diameter less than 1 micrometer were eliminated for the following analysis. The morphological metrics, including fiber diameter, myelin thickness, axon diameter, and g-ratio (ratio between axon diameter and fiber diameter) of sampled myelinated fibers were extracted by using AxonDeepSeg software (Figure S1B)^93^. The conduction velocity of quantified fibers was estimated by simulating the multi-compartment axon model of Halter and Clark^94^ implemented in the myelinated axon model^18,55^.

**Figure S1:**
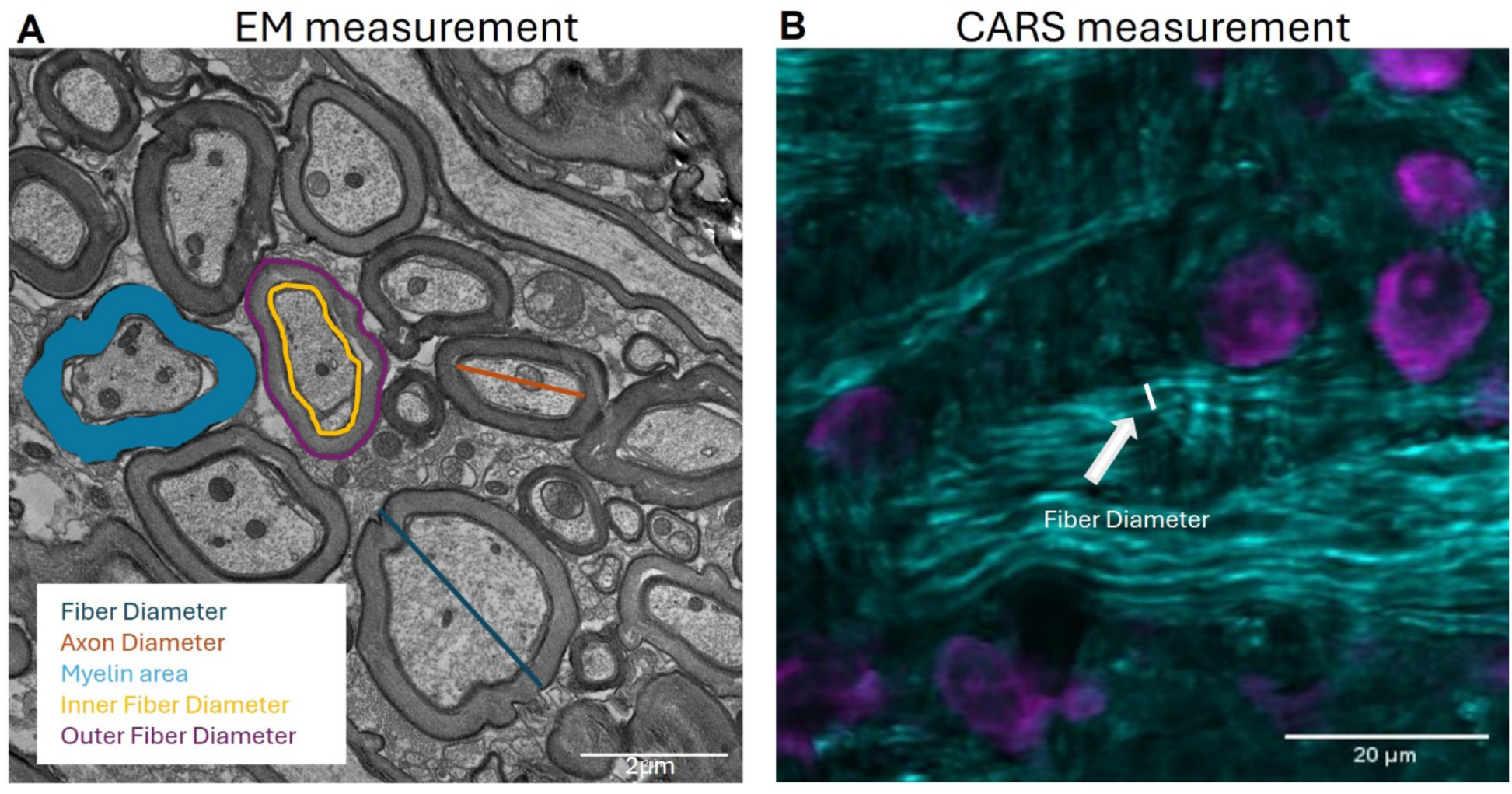
Quantification of myelination morphology in CARS and EM images. EM images provide higher resolution for detailed analysis, including fiber diameter (dark blue), axon diameter (teal), myelin thickness (green), inner (orange), and outer (purple) fiber diameter **(A).** CARS images were quantified using the FIJI line segment tool; a white line **(B)** was drawn, and the measure function was used to determine fiber diameter for 20 axons per image (Nissl-stained cell bodies in purple, CARS signal in cyan).

### Pre-pulse inhibition of the acoustic startle

The methods for prepulse inhibition followed the groups previous publication by Greene et al.^48^.

Apparatus: All experiments were conducted in a double-walled, sound-attenuating chamber (interior dimensions: ∼3 x 3 x 3 m; IAC, Bronx, NY) lined with echo-attenuating acoustical foam. Animals were placed in an acoustically transparent cage constructed of aluminum wire mounted to a flexible polycarbonate platform and raised to speaker height by four polycarbonate (PVC) posts (Figure 7A). The aluminum wire cage was constructed to allow for enough animal movement to generate startle responses, but not enough room for the animal to turn its head or turn around in the cage. This design allowed us to keep the animals positioned at a midline speaker during spatial speaker swap tasks. All animals were tested in the dark and were visually monitored using a closed-circuit infrared (IR) camera to ensure the correct orientation. Prepulse stimuli were presented from 25 loudspeakers (Morel MDT-20) spaced along a 1 m radius semicircular boom at 7.5° increments, from 90° (right) to 90° (left). The speaker boom was oriented horizontally (i.e. 0°elevation) for all experiments. Startle-eliciting stimuli were presented from a Faital Pro HF102 compression driver mounted ∼25 cm above the animal and amplified with a Yamaha M-40 power amplifier. The startle response of the animal was captured using a cage-mounted accelerometer (Analog Devices ADXL335).

Three Tucker-Davis Technologies (TDT) RP2.1 Real-time Processors controlled by custom written MATLAB (MathWorks) software generated the stimuli and recorded the startle response. Startle-eliciting stimuli (SES) were 20ms duration uniformly distributed broad-band noise bursts (rectangular-gated, 50 kHz bandwidth) dynamically generated by the first RP2.1, and presented at 117 dB SPL, except when testing startle response of gerbils to different startle speaker levels (Figure 7B, 8C). Carrier stimuli (CS) consisted of uniformly distributed broad-band noise dynamically generated by the second RP2.1 and presented continuously (except when otherwise noted) during testing. In some experiments the broad-band noise CS was low-, high-, or band-pass filtered with a 100th order FIR filter that was designed in MATLAB and implemented in the second RP2.1. The CS was presented from one speaker at a time, controlled by switching the output from channel 1 to 2 on the third RP2.1 (which received the CS as an input). The speakers corresponding to these outputs were dynamically set by two sets of TDT PM2Relay power multiplexers, controlled by the first and third RP2.1. The motion of the polycarbonate plate resulting from the startle response was transduced by an accelerometer, and the voltage output sampled at 1kHz by the first RP2.1. The startle response amplitude was calculated as the RMS of the accelerometer output in the first 100 ms period after the delivery of the SES. Prepulse and startle stimuli presentation was identical in setup to previously run experiments by the lab ^44^.

Prepulse Inhibition: Conditions were presented 11 times in all experiments, with the first ten conditions being used to calculate PPI value. Animals were limited to <60 mins of testing each day to avoid habituation to the startle stimuli. In experiments with more than 10 repetitions recorded, testing was spread out over multiple days. Gerbils were given at least one recovery day in between testing days to further avoid habituation to the startle stimuli. Experimental and control trials within each testing day were presented pseudo randomly. Stimulus presentation and response measurement was computer controlled. For each prepulse task, startle amplitudes from each repetition were averaged for every condition. In tasks where multiple days of testing were performed, reptations spanning multiple days were averaged together for each condition. Startle amplitude from control conditions (no prepulse cue presented) were averaged together as well. The PPI value for each condition was then calculated as 1-(Startle in Prepulse conditions/Startle in Control Condition). Since PPI value reflects a reduction of startle amplitude, a value closer to 1 means higher suppression of the startle response, and therefore better detection of the prepulse, while a value closer to 0 means lower suppression of the startle response, and poorer detection of the prepulse. To quantify this, if an ANOVA revealed a condition that was statistically higher than 0, that would indicate significant detection of the prepulse in that condition, while one that is not statistically above 0 would indicate no significant detection of the prepulse.

Startle Amplitude Response: Gerbils were held in the apparatus described above at the midline (0°) speaker. Instead of presenting the startle stimuli at 117 dB, as for all prepulse experiments, we varied the decibel level of the startle stimuli between 80 and 130 dB in 5 dB level steps. Each day of testing lasted approximately 45mins. The startle amplitude of gerbils was measured in response to the different level presentations of startle. 11 repetitions of each condition were run, with 10 being measured and recorded. The session started with a two-minute acclimation period of no startle, and the inter-trial interval (ITI) was averaged at 20s, which varied between 15 and 25s at an interval of 1s. The startle amplitude response was measured in the presence of 70dB background broadband noise from our speaker apparatus.

Gap Detection: We used gap detection to test the ability of gerbils to utilize a prepulse inhibition cue as a metric of detection, and also to test the temporal fidelity of gerbils to a cue of varying length. Each condition was repeated 11 times, with 10 conditions being recorded. Each day of testing lasted approximately 30 mins. In addition to the 7 experimental conditions, 2 control conditions with no prepulse were intermixed in the testing. The apparatus, acclimation period, and ITI of conditions were identical to the startle response tests listed above. In the gap detection tasks, and in all following prepulse tasks, startle speaker SPL was consistently 117 dB. 70 dB broadband noise was presented from our speaker array. The prepulse in the gap detection task was a gap in broadband noise of variable length of either 1, 3, 5, 10, 15 or 20ms, which was constantly presented 80ms prior to startle presentation.

Minimum audible angle: We used the speaker swap ^45^ paradigm to assess minimum audible angle in our gerbils. Each test session lasted <40 mins. To test responses across the animal midline, we held animals towards the 0° center speaker in the apparatus described above. Startle SPL, acclimation period, and ITI of prepulse stimuli were identical to experiments listed above. The prepulse in SSwap experiments was a change in the presentation location of the CS between two matched speakers with various separation angles. Prepulses were presented 80 ms prior to startle speaker presentation in all swap angle conditions, as this was sufficient time to elicit inhibition of startle in gerbils, but not so long as to lose the inhibition effect of the prepulse. We ran 30 repetitions of each experimental (angle of speaker swap) condition, which was spread out over three days of testing at 10 repetitions each day.

### Statistical Analysis

For CARS and confocal images, we employed ANOVA followed by the Tukey test to measure statistical differences between MNTB regions in young and aging groups. Student’s t-test was used to measure statistical differences in each MNTB subregion between the two age groups. Reported p values were corrected for multiple comparisons using the Bonferroni correction. For metrics quantified from ABRs and EM images, the two-sided Mann-Whiteny U test was utilized due to its robustness in handling uneven sample sizes and skewed distributions.

For behavior experiments, statistical analysis was performed using GraphPad Prism. Normal distribution was assumed for all data. For each task, a two-way ANOVA was performed, with the main effect factors being experimental condition (dB level for startle amplitude test, gap duration for gap detection test, speaker swap angle for speaker swap test) and age group. Post-hoc comparisons were assessed in each ANOVA to compare conditions both across age group and within age group. Significant detection was determined if the PPI level for an experimental condition in each age group was significantly above 0.

